# Classification and Generation of Microscopy Images with Plasmodium Falciparum via Artificial Neural Networks

**DOI:** 10.1101/2020.07.21.214742

**Authors:** Rija Tonny Christian Ramarolahy, Esther Opoku Gyasi, Alessandro Crimi

**Affiliations:** African Institute for Mathematical Sciences, Summerhill Estates, East Legon Hills, Santoe, Accra, Ghana; Marine Ecology Department, Institute of Marine Sciences Kiel, P.O. Box LG 581 Legon, Accra, Ghana

**Keywords:** blood smear, machine learning, deep-learning, convolutional neural network (CNN), image processing, automated detection, generative adversial networks (GANs)

## Abstract

**Background:** Recent studies use machine-learning techniques to detect parasites in microscopy images automatically. However, these tools are trained and tested in specific datasets. Indeed, even if over-fitting is avoided during the improvements of computer vision applications, large differences are expected. Differences might be related to settings of camera (exposure, white balance settings, etc) and different blood film slides preparation. Moreover, generative adversial networks offer new opportunities in microscopy: data homogenization, and increase of images in case of imbalanced or small sample size.

**Methods:** Taking into consideration all those aspects, in this paper, we describe a more complete view including both detection and generating synthetic images: i) an automated detection used to detect malaria parasites on stained blood smear images using machine learning techniques testing several datasets. ii) investigate transfer learning and further testing in different unseen datasets having different staining, microscope, resolution, etc. iii) a generative approach to create synthetic images which can deceive experts.

**Results:** The tested architecture achieved 0.98 and 0.95 area under the ROC curve in classifying images with respectively thin and thick smear. Moreover, the generated images proved to be very similar to the original and difficult to be distinguished by an expert microscopist, which identified correcly the real data for one dataset but had 50% misclassification for another dataset of images.

**Conclusion:** The proposed deep-learning architecture performed well on a classification task for malaria parasites classification. The automated detection for malaria can help the technician to reduce their work and do not need any presence of experts. Moreover, generative networks can also be applied to blood smear images to generate useful images for microscopists. Opening new ways to data augmentation, translation and homogenization.

## Background

Malaria is an infective disease caused by a blood parasite of the genus Plasmodium. According to the latest World malaria report, there were 228 million cases of malaria in 2018 compared to 231 million cases in 2017 [1]. Five different types of parasites exist: Plasmodium falciparum, Plasmodium vivax, Plasmodium malariae, Plasmodium ovale and Plasmodium knowlesi. The plasmodium falciparum is the most lethal and accounts for almost 99% of the malaria cases in sub-Sahara Africa. Besides, each parasite has a distinct form depending on its development. Indeed, parasites go through different developing stages: ring stage, trophozoide stage, schizont stage and gametocyte stage. These stages help parasitologists to know the types of parasites as they differ from each other [2]. **Microscopy** investigation remains the major form of diagnosis in malaria management in low- and middle-income countries. Although, it has been shown that imprecision might occur due to non-expert microscopists especially in rural areas where malaria is endemic [3].

Other methods include **rapid diagnostic tests (RDTs)**, which are easier to use but have lower sensitivity and specificity than microscopy approaches, and it has been estimated that they are not cost-efficient in some geographical locations [2]. **Polymerase chain reaction (PCR)** gives a better result than the two previous methods, having more sensitivity and specificity [4]. However, it is limited by the need of experienced user, reagents, and expensive devices [2, 5, 4]. In summary, for low- and middle-income countries, microscopy is still the most practical choice for malaria diagnosis.

Microscopy diagnosis initially requires determining the presence of malarial parasites in the specimen in exam. Then, if parasites are detected, species identification and calculation of the degree of infection (number of parasites) is performed. Those tasks are currently still mostly carried out by a human operator, leading to variability and possible imprecision [3]. Recently, advancements in computer vision are allowing malaria diagnosis in microscopy in an automated manner by using image processing and machine learning. The image processing step has the purpose of detecting/segmenting the region of interest (ROI) inside blood smears from a microscope by using image processing techniques. The ROI selection is generally performed by Otsu thresholding [6]. Given the high level of overlapping and touching red blood cells in the thin smear staining further processing is needed. Rajaraman et al. [7] used a multi-scale Laplacian of Gaussian filter as an edge detector, and a level set activation contour technique as segmentation technique. Other works used the watershed approach to separate overlapping/touching red cells [8, 9]. ROI detection is more convenient than full sliding window at different scales due to computational costs [9]. The second step is given by a classifier which discriminates whether the detected elements in the ROI is an infected red blood cell or not. Before the deep-learning era, classifiers such as *support vector machine (SVM)* [9], *logistic regression* [10], *Bayesian learning* [9] or other machine learning techniques were used, and an extra step for feature extraction and selection were needed. This feature selection required an engineering expertise to define a specific feature to be extracted as color, textural and morphological features to distinguish the parasitized and healthy cells [2]. Furthermore, machine learning algorithms have been used to go beyond mere detection but also to differentiate among the parasites. For instance, Das et al. used Näive Bayes algorithm to classify ring and gametocyte stages of P.vivax and P.falciparum achieving high accuracy [9]. Anggraini et al. classified the malaria parasites of different stages such as ring, gametocyte and other artifacts by using Bayesian classifier [11]. More recently, by using deep-learning models such as *convolutional neural network (CNN)*, the step of hand-engineered feature selection is not needed, CNN training performs feature extraction jointly to the training. Moreover, as in many field, deep-learning methods outperform other machine learning methods [12]. Also, deep-learning architectures introduced the concept of transfer learning, where pre-trained models are either fine-tuned on the underlying data or used as feature extractors to aid in visual recognition tasks [13]. These models transfer their knowledge gained while learning generic features from large-scale datasets. The transfer of previously-learned skills to a new situation is generalized, rather than unique to the situation. Rajaraman et al. investigated how known pre-trained models (AlexNet, Visual Geometry Group architecture 2016 (VGG-16), ResNet-50, Xception, DenseNet-121) perform transfer learning on malaria images [7]. Moreover, neural networks diagnoses have been compared to results obtained by using real-time PCR, and human analysis of microscopy images to obtain comparable results in an in-field study in Peru [14]. In this study the architecture was a VGG-16 trained for the purpose [15]. Lastly, some studies included these image processing approaches in mobile phone applications with the goal of making the analysis accessible in remote settings without access to computers [6, 16].

When using a machine learning based classifier, the number of available samples for training is an important key to high accuracy. However, obtaining images may be expensive and time consuming, and often the obtained image datasets are imbalanced. Namely, many sample of one specific plasmoid or uninfected are available and very few of a specific instance are available [17]. In this context and in other computational pathology contexts, data augmentation is rising as a promising solution. A computational architecture called *Generative Adversial Networks (GANs)* allows the generation of synthetic images from very small training images available [18]. In this manner, deep-learning architectures are not uniquely used to detect or classify, but they can also create new data. GANs architectures are composed by two sub-systems which compete with each other: a *generator* and a *discriminator*. They can be seen as crook printing *fake money* and a bank clerk verifying *money*, the first trying to create the most realistic fake money and the latter becoming better at distinguishing fake from real money. In our case, money is a metaphor representing microscopy images. GANs architectures have reached incredible level of accuracy with humans not being able to distinguish the real by the generated images [19]. The use of GANs in pathology is motivated by major factors related to data augmentation: 1) to offer variability to a pathologist learning possible variation of the images in study. 2) to have more balanced data (as usually control images are more available than specific case images). 3) data translation and homogenization. Therefore, a complete deep-learning tool should include both discriminator and a generator part to benefit also the discriminator [20]. Therefore, we present here a comprehensive analysis on malaria images both in terms of automated classification and generation for thick and thin smear.

## Methods and Materials

### Data and Code

In this paper, we used three different datasets with films stained according to the Giemsa protocol and all focused on Plasmodium falciparum. The three datasets are publicly available. All data have been collected according to ethical standards following the declaration of Helsinki, informed consent was obtained from all the subjects. Ethical approval has been previously granted by local committees for the individual studies though they are independent from the study reported in this paper.

- A cell dataset containing 27, 558 individual images of red blood cells, taken by a smartphone mounted on a microscope at Chittagong Medical College Hospital, Bangladesh [7]. The images comprise equal instances of infected and uninfected cells. The individual cell images are taken from stained thin blood smears of 150 infected and 50 healthy patients. The images were manually annotated by an expert slide reader at the Mahidol-Oxford Tropical Medicine Research Unit in Bangkok, Thailand. The dataset is accessible at the URL https://lhncbc.nlm.nih.gov/publication/pub9932.
- A dataset including 1182 pictures of stained thick blood smear taken from a smartphone attached to a microscope all presenting at least 1 infected cell, using field stain at x1000 magnification. The images were acquired and annotated by experts of the Makerere University, Uganda [16]. The dataset is accessible at the URL http://air.ug/downloads/plasmodium-phonecamera.zip.
- A dataset comprising 655 images of thin smears acquired with a magnification of 100*X* by an immersion objective of Leica DLMA microscope [21], all with at least 1 infected cell. The dataset is publicly available at the URL https://drive.google.com/open?id=1EMJ7dg0TBs34sDWcj7Tj1wozXJC0wtbc The produced software tools are publicly available at the URL https://github.com/alecrimi/malaria_detection_and_generation.

### Parasites detection

The detection experiments are performed on a trained CNN models which are fed with uniformly scaled patches of candidate regions potentially containing parasites. In this work we also adopt a ROI detection approach to identify the patches which are fed into the CNN as done in most of the aforementioned studies. More specifically, except the single cell dataset, we use the Otsu automatic thresholding to separate the image into background and foreground. Then, we remove the noise using the morphological operation opening and increase the area of foreground using dilation. Finally, we apply the watershed algorithm to mark each border of each object using the label from the connected components. Candidate ROIs to be checked are scaled to 100×100 pixels for consistency to the classifier. The used CNN model is inspired by the LeNet model [22], and it comprises of three hidden convolutional layers. *Convolutional layers* are those effectively extracting autonomously the features from image. Then, inspired by biological neurons each convolutional layer is followed by an activation function which emulates the spike of neurons (in our model *rectified linear units (ReLu)* are used). *Pooling layers* are used to take the dominants features given by the convolutional layers and they are also used to reduce the dimensionality of the parameters of the image. Fully connected layers are computational units driving the final classification decision. Drop-out is the procedure of removing randomly some connections between layers. Dropping out some neurons which have less probability will give a best result and make the training faster. The last layer is the one responsible of classification. In the reported experiments, it was trained to discriminate between image containing parasites or not, though it can be easily expanded to several classes (ring stage, trophozoide stage, schizont stage and gametocyte). In this study, the goal is limited to detect and count any form of parasites. The whole architecture with its parameters is summarized in Table 1 and Figure 2. More specifically, the input image for the first dataset has a pixel resolution 100 *×* 100 *×* 3 and 50 *×* 50 *×* 3 for the second dataset. The first convolutional layer is made of 32 kernels, the second is made of 64 kernels and the third has 128 kernels; all with the same size 3 *×* 3. ReLu is applied to the output of each convolutional layer as the activation function and each convolutional layer is followed by a max pooling of size 2 *×* 2. The output of the last max pooling is flatten into a vector and becomes the input of the first fully-connected layer. The fully-connected layers have 512 neurons each and ReLu is also applied to their outputs as the activation function. Each fully-connected layer is followed by a dropout with probability of 0.3 and the output of the last dropout feeds into a sigmoid classifier. These layers are also depicted in Figure 2 where each block represents a convolutional filters bank or a fully connected layer. The pyramids represent the maxpooling downsampling. The inner parallelepipeds represent the size of the convolution filter. Last layer is of size 1 *×* 1 as the sigmoid functions have to decide whether there is a parasite in the ROI in exam or not, though it can be expanded into more classes to discriminiate the development stage or plasmodium type.

**Table 1.** Used CNN model and its parameters

| Type | Kernels/Neurons | Shape (size) |
| --- | --- | --- |
| Input | | $100 \times 100 / 50 \times 50$ |
| Conv1 + ReLu | 32 | $3 \times 3$ |
| Maxpooling1 | | $2 \times 2$ |
| Conv2 + ReLu | 64 | $3 \times 3$ |
| Maxpooling2 | | $2 \times 2$ |
| Conv3 + ReLu | 128 | $3 \times 3$ |
| Maxpooling3 | | $2 \times 2$ |
| Flatten |  |  |
| FC1 + ReLu | 512 |  |
| Droupout (0.3) |  |  |
| FC2 + ReLu | 512 |  |
| Droupout (0.3) + Sigmoid |  |  |

**Figure 1.**
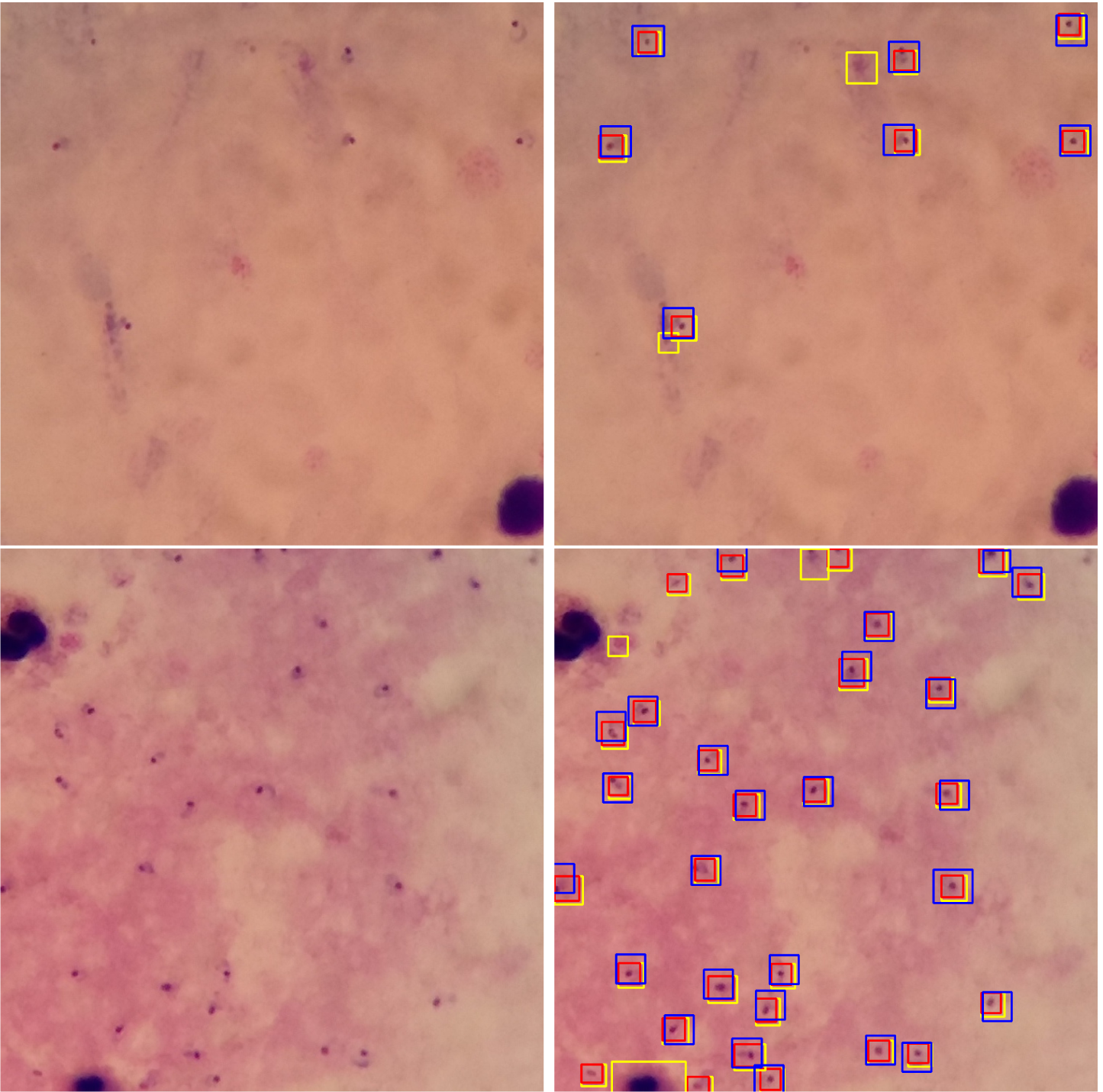
Parasites detection on a thick smear image. On the left is the original image and on the right, the parasite detection. The yellow rectangles show the candidate parasites, the red boxes show the detected parasites from the model and the blue boxes show the ground-truth parasites annotated by a microscopist. The examples on the bottom row shows a white cell initially selected as a ROI but then discarded by the classifier. In this latter image, two false positive -according to the microscopist-detection are also present.

**Figure 2.**
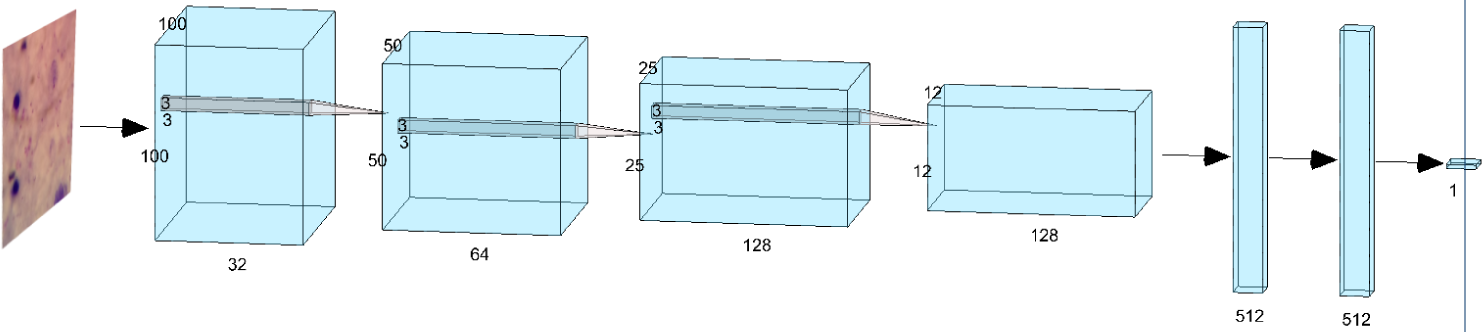
CNN architecture for classification; each block represents a convolutional filters bank or a fully connected layer. The pyramids represent the maxpooling downsampling. The inner parallelepipeds represent the size of the convolution filter.

Plasmodium species are distinguishable from regular blood components and artefacts by their characteristic shapes and color properties [23]. Giemsa-stained microscope images provide good contrast between deep purple nuclei and background. Hence, both in classification and generation, color information is used. This might lead to higher number of used parameters though in parallel aids the classifation and makes image generation more interesting. Therefore, both classification and image generation use images having the 3 color channels.

In the following section we report the results using the trained CNN for the thick and thin smear dataset. Moreover, we further test the trained thin smear model on an unseen dataset, and we repeat the experiments by carrying out transfer learning on a VGG architecture as used in previous literature [7, 15].

### Image generation

For the image generation, deep convolutional networks are also used, but as mentioned earlier, the architecture is more complex and divided into generator and discriminator as shown in Figure 3, rather than just a discriminator. Being two sub-systems, they can have independent sets of hyper-parameters. A relevant aspect is that the generator should have a faster convergence than the discriminator, otherwise the discriminator might always understand that the images are fake hence not allowing proper image generation. A solution is to give smaller learning rate. In our experiments, learning rates of 0.00001 and 0.0002 were respectively used for the discriminator and the generator. Moreover, 1000 iterations were used to allow the generator to create realistic images with an output size of 256×256 pixels. A further relevant detail is represented by how the expansion steps are carried out, as the typical checkboard artifacts can appear if deconvolution is used as an expansion. The adopted solution is to simply switch out the standard deconvolutional layers for nearest-neighbor resize followed by convolution causing artifacts of different frequencies to disappear [24]. The used GAN architecture was designed inspired by the suggested structure in [25] as depicted in Figure 3. Given the fact that each image has several elements in it, and the desire of simulating a circumstance with few samples available, the image generation is performed by using 10 randomly selected images from the available dataset. For the single cell and thick smear dataset, images are used to train as untreated, while for the thin smear dataset images were also contrast adjusted as it was noticed to improve the results. To assess the quality of the generated images, an experienced microscopist has been involved, being unaware of the process and of the number of generated images, only aware that some images were synthetic and some real. The micscopist received jointly all images. More specifically, he received 10 images comprising single infected cells, 6 images of thick staining and 4 images of thin staining.

**Figure 3.**
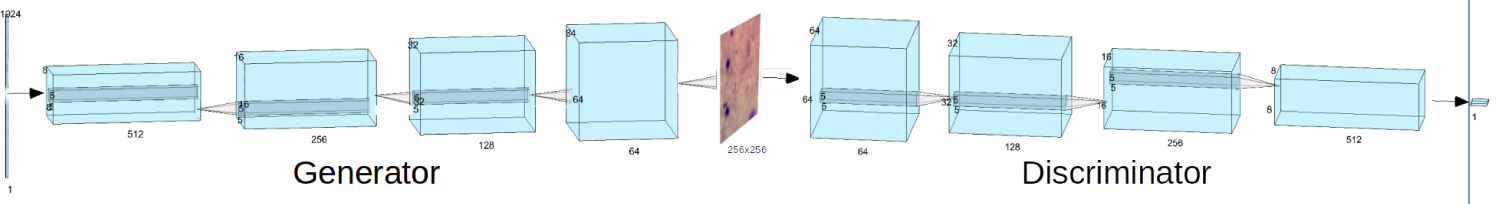
GAN architecture, each block represent a convolutional filters bank or a fully connected layer. The pyramids represent the either the upsampling or downsampling. The inner parallelepiped represent the size of the convolution filter.

## Results

### Parasites detection

The parasite detection achieved satisfactory results in line with previous literature with an AUC is equal to 0.98 and 0.95 respectively for the thick and thin staining. The performance metric for the models trained and tested on the two datasets are shown in the Table 2. It is worthwile to mention that sometimes the selected ROIs were containing white blood cells which were then discarded by the neural networks as infected.

**Table 2.** First 2 rows are the performance of the model described in Table 1. Third row reports the performance of the mixed model: CNN described in Table 1 trained on the single cell dataset and tested on the thin smear dataset.

| | Accuracy | Precision | Recall | $F_1$ score | Sensitivity | Specificity |
| --- | --- | --- | --- | --- | --- | --- |
| Individual cells | 0.953 | 0.953 | 0.953 | 0.953 | 0.947 | 0.959 |
| Thick patches | 0.989 | 0.989 | 0.989 | 0.989 | 0.993 | 0.982 |
| Mixed model experiment | 0.895 | 0.893 | 0.895 | 0.895 | 0.894 | 0.896 |

As shown in Table 2, our models performed a satisfactory classification for the two datasets. For the first model, the accuracy value shows that 95% of the first dataset were correctly classified and for the second model, 98% of the second dataset were correctly classified. The high values of the precision, recall and *F*1 score for the two models give information about their performances. Despite the difference in size between the patches and the input of the model, Table 2 also shows acceptable performance of the mixed test because from the accuracy value, 89% of the second dataset were correctly classified by the first model. And also, the value of precision, recall and *F*1 score are high. The proposed model outperformed the VGG architecture especially in the mixed model experiment [7, 15].

Figure 1 shows the detection of parasites in a thick blood smear image. The first figure is the original image and the second is the result of our algorithm. The yellow rectangles show the candidate parasites, the red boxes show the detected parasites from the model and the blue boxes show the parasites from the annotations data. Figure 4 shows the result of the segmentation using watershed transform and detection of the infected cells. The first figure (left) is the original thin smear image, the second figure (right) shows the result of our segmentation which gives satisfactory results and the third figure gives the infected cells predict by our model. The cells marked in red are those detected as infected by the model.

**Figure 4.**
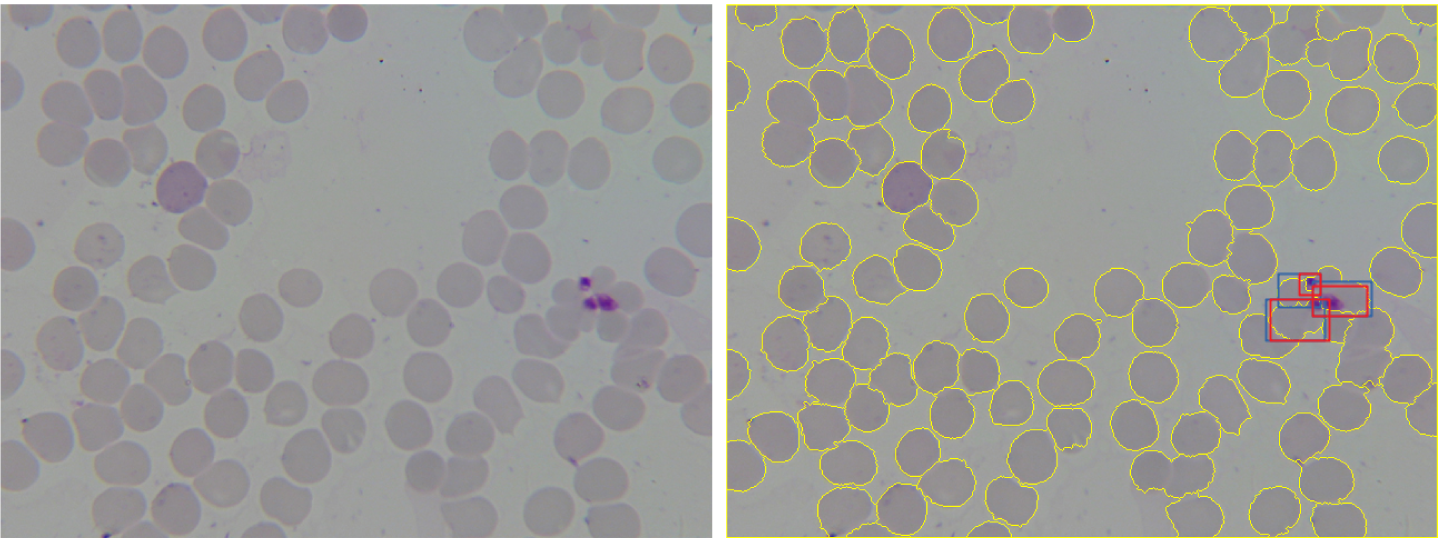
Cells segmentation and infected cells detection on a thin smear image. On the left original image, on the right the results of the detection. The yellow lines show the candidate parasites, the red boxes show the detected parasites from the model and the blue boxes show the ground-truth parasites annotated by a microscopist.

The training was performed on a Google Colab environment using 1x GPU Tesla K80, having 2496 CUDA cores, and 12GB GDDR5 VRAM; and 1 single core CPU hyper threaded Xeon Processors 2.3Ghz and 12.6 GB RAM available. The training time was 2.41 minutes, while the testing time was less than a second per image.

### Image generation

The computational time to produce a single image was 3 hours on a single node cluster on a Openstack based system with 125GB RAM and 32 VGPU and CUDA enabled. The resulting images given to the microscopist during the test are reported in Figure 5, 6 and 7. It was noticed that with the used GANs architecture, the thin smear image generation was not as satisfactory than the other datasets.

**Figure 5.**
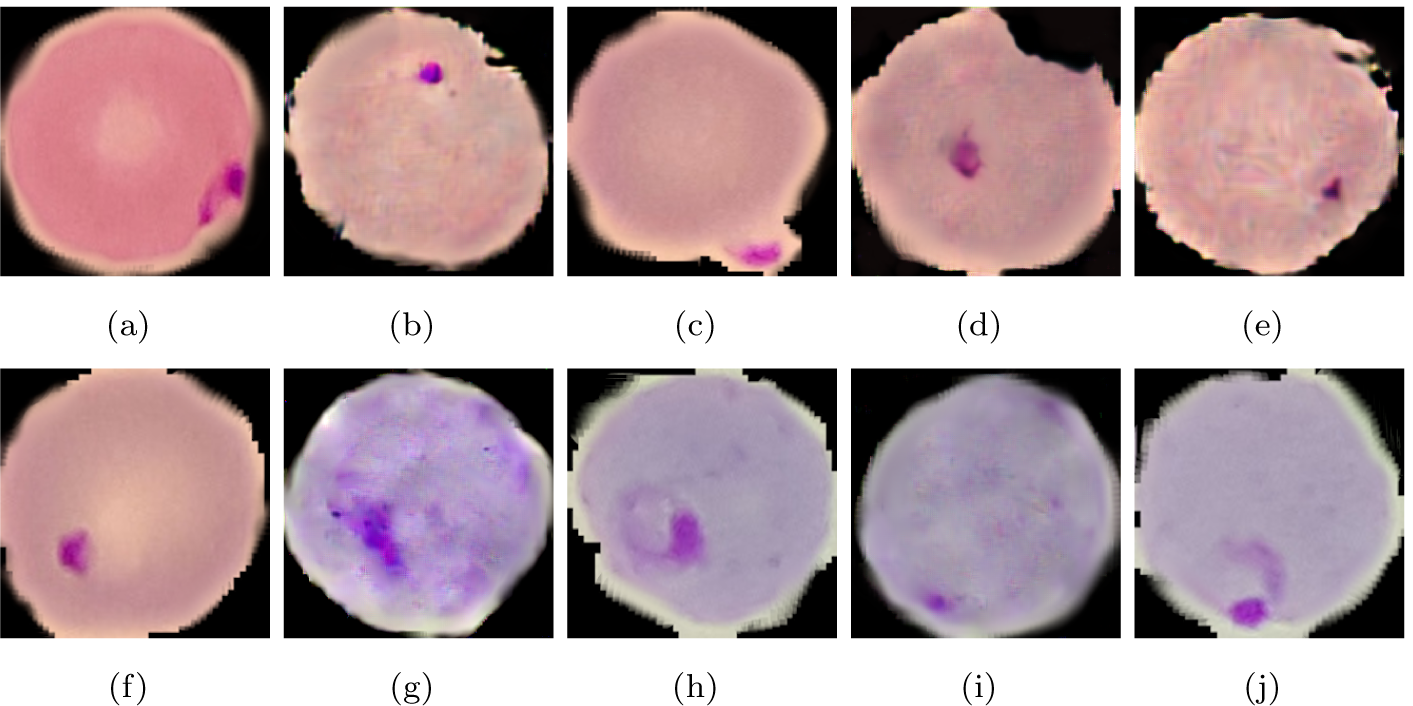
Single cells infected by parasite showing different staining. Real: f, h, j. Synthetic: a, b, c, d, e, g, i.

**Figure 6.**
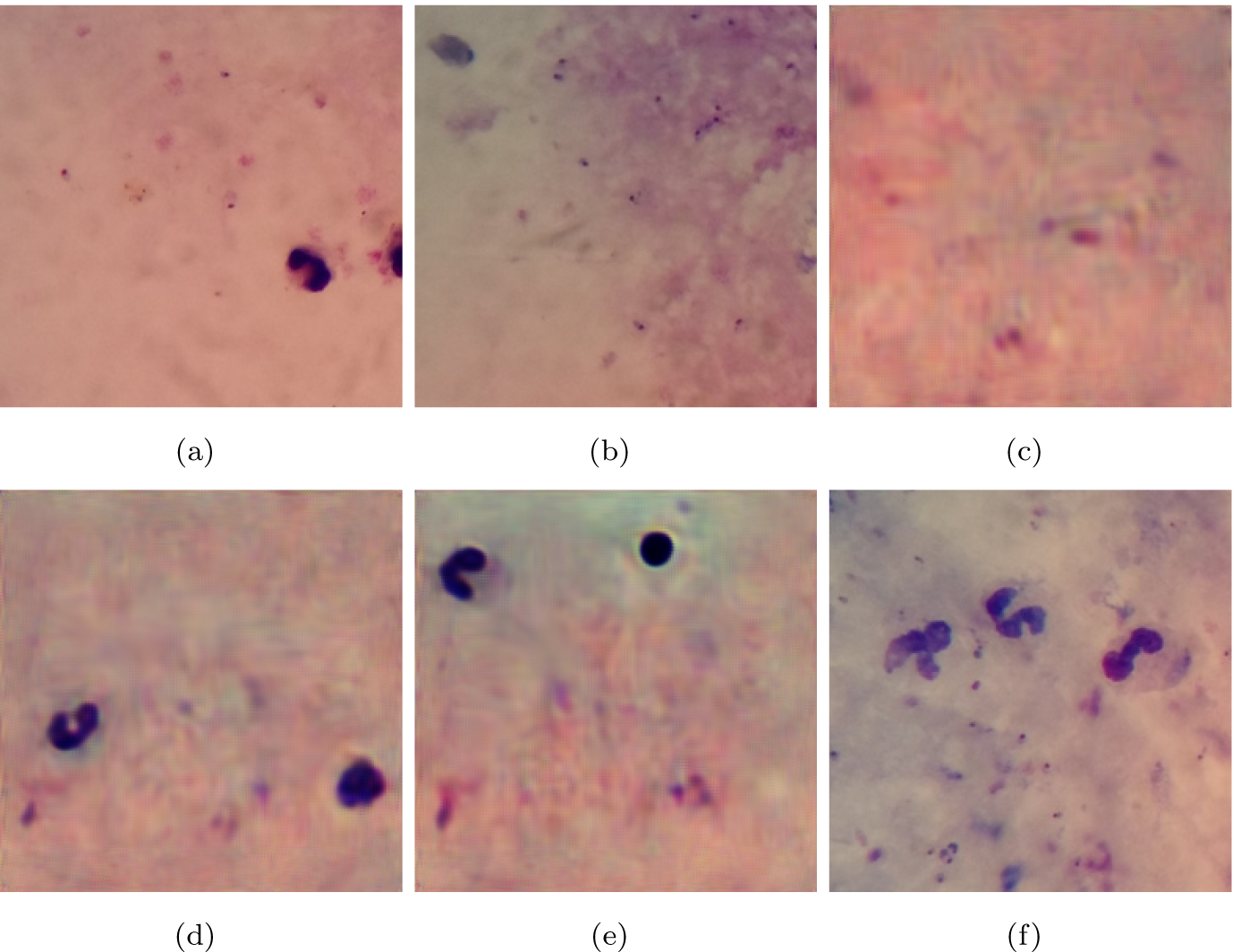
Thick staining images. Real: a, b, and f. Synthetic: c, d, and e. In those images, it seems visible that GANs can easily generate white blood cells given their size, but has some challenges in creating sharp edges for the smaller red blood cells.

**Figure 7.**
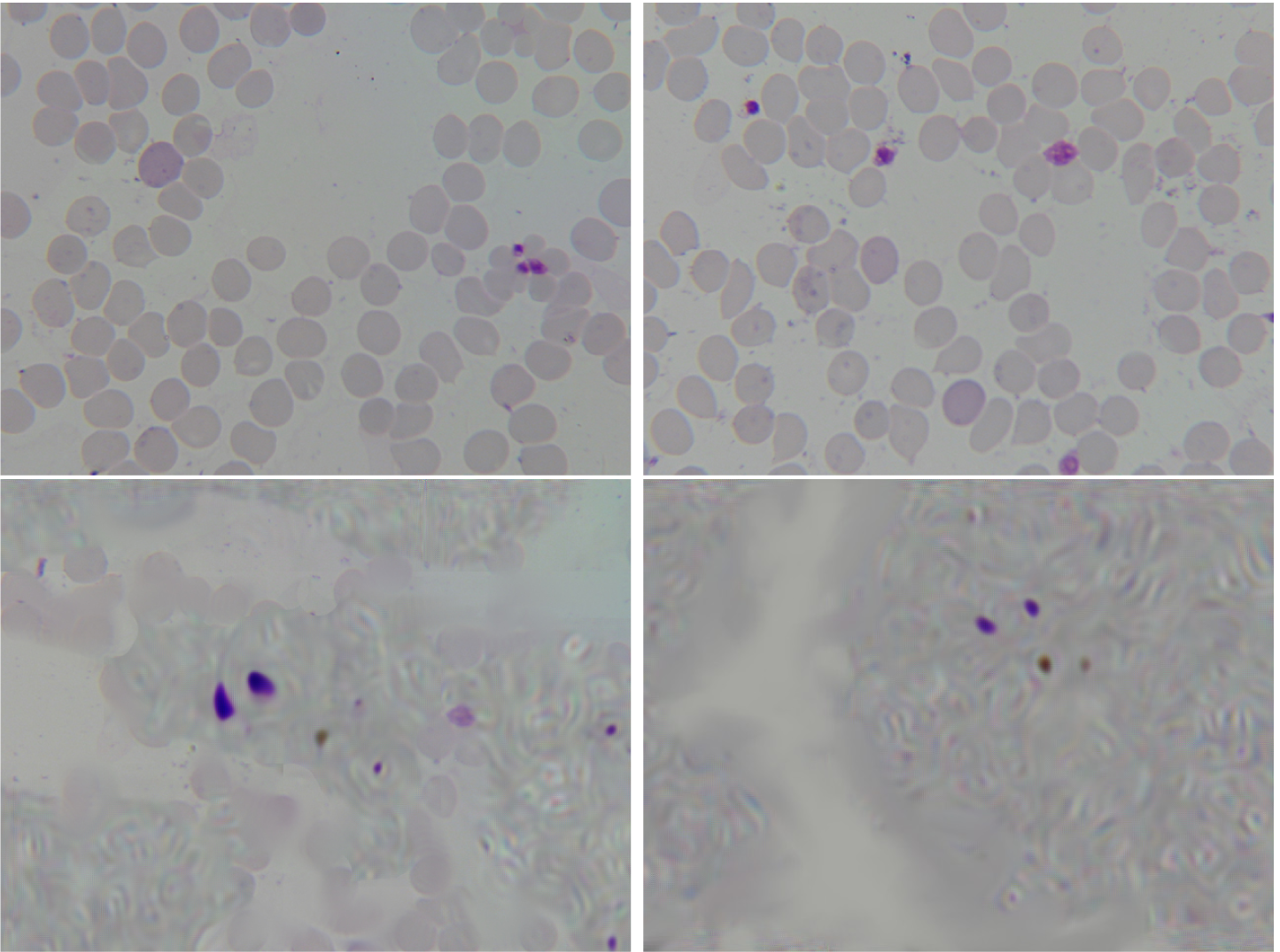
Thin smear images given to the microscopist to guess which real and which synthetic. Top row real images, bottom rows synthetic images.

**Table 3.** First 2 rows are performance of transfer learning using a pre-trained VGG-19. Third row reports the performance of the mixed model using the VGG-19, trained on the single cell dataset and tested on the thin smear dataset.

| | Accuracy | Precision | Recall | $F_1$ score | Sensitivity | Specificity |
| --- | --- | --- | --- | --- | --- | --- |
| Individual cells | 0.940 | 0.940 | 0.940 | 0.940 | 0.932 | 0.947 |
| Thick patches | 0.985 | 0.985 | 0.985 | 0.985 | 0.987 | 0.982 |
| Mixed model experiment | 0.697 | 0.767 | 0.697 | 0.620 | 0.682 | 0.920 |

The microscopist (EKN) identified correctly the real images with thick staining, but misclassified 5 out of 10 images in the single cell test. Moreover, the microscopist described having difficulties in taking a decision with the image depicted in Figure 6 (a). From the thin smear images, the microscopist had only one mistake. Observing, the generation of images through the iterations, it can be seen that the first 300 iterations produce meaningless blobs, and only around 700 iterations the generated images appear meaningful. Moreover, the experiment confirmed the need of using slower convergence of the discriminator compared to the generator. If this approach was not used, the generator was never going to be able to produce plausible images.

## Discussions

Detection of malaria infected erythrocytes from peripheral blood smear samples using light microscopy is a challenging task for humans, so it is for software tools given the presence of artifacts. In this study we investigated both classification of plasmodium in microscopy images, as well as generation of new data for data augmentation.

We considered microscopy images stained by using the most common Giemsa staining. More specifically, thick and thin smears containing Plasmodium falciparum previously annotated by experts. In contrast to legacy machine learning models which need further feature extractions, deep-learning models require less work and achieve more accuracy. Dong et al. [12] reached an accuracy of 0.92 using SVM, and Das et al. [9] got an accuracy of 0.84 and 0.83 with Bayesian learning and SVM respectively for malaria parasites classification. The proposed deep-learning approach gave satisfactory results with AUCs of 0.98 and 0.95 in a cross-validation setting. In the used CNN architecture inspired by LeNet [22], implicit regularization imposed by batch normalization, and dropouts in the convolutional and dense layers led to improved generalization. To be in line with previous research using deep-learning on microscopy images we used the a pre-selection of ROIs and then evaluated them [21, 10, 9, 26]. Nevertheless, recent architectures tested in other object detection task can also perform detection while performing classification, in this way avoiding the initial step of ROI detection. The most successful technique is called *You-Only-Look-Once* (YOLO) [27]. Given the excellent results obtained with the proposed approach, we consider simultaneous detection/classification via CNN as a future improvement. The transfer learning experiment gave also very promising results, with the VGG architecture having almost the same accuracy of our proposed CNN model.

Automated detection for malaria can help the technician to reduce their workload. Indeed it has been reported that a microscopist need an average of 50 minutes to evaluate a smear [28], while given the need of few seconds to detect infected cells with the proposed tools, the time can be reduced to few minutes plus the time of the staining. The algorithm focuses mostly on detection of infected cells, as the count of those cells can give an indication of the level of infection. Indeed, the number of cells needed to diagnose is defined as the number of red blood cells to be tested to yield a correct positive test [7].

Moreover, in this article, it was also shown that GANs can be used to generate synthetic infected blood smears which look realistic, and can be used to increase human training on how differently a pathogen can appear and for data augmentation to further data analysis. To our knowledge, this is the first study generating synthetic microscopy images with malaria parasites. The experienced microscopist involved in the study, reported anecdotally that the discrimination between real and synthetic images was challenging. Without context, he had 50% of misclassification with the single cells images. Instead, he successfully identified all real and synthetic images for the thick smear. A possible explanation is given by the fact that the GANs were able to generate plausible white blood cells and artifacts, but the small erythrocytes were blurred in the synthetic images and sharper in the real one, and this led the microscopist to identify correctly the images. Regarding the thin smear images, it was noted that the synthetic images had several ringing artefacts, this is probably related to extreme upsampling. We gave only 4 images to the microscopist which only misclassified one, we limited to 4 as the generated synthetic images were not satisfactory as for the other datasets. We speculate that we have been able to deceive the microscopist due to the fact he had only four examples to look at, and with more data he would have been able to guess better patterns noticing the presence of ringing artifacts in some images rather than others. Nevertheless, data augmentation has become common tools in several digital pathology context [29], so it will be for malaria studies. Moreover, GANs are often used to translate data from one domain to another, e.g. to move from a magnetic resonance scan to a dyed histopatological images and the other way round [30]. There has been a recent growing interest in applying GANs to solve several tasks related to digital pathology, including staining normalization [31] and staining transformation [32, 33]. In this first study on microscopy images for malaria, we showed that it is in the first place possible to generate realistic synthetic images for malaria analysis. This demonstration can inspire researchers. For example, one can uniform slides acquired in different centers into homogeneous slide style or other generative results for digital pathology.

## Conclusions

In this study we proposed a neural network architecture able to classify infected cells in microscopy images with high accuracy and in different datasets. This shows the potential of deep-learning architecture to be adopted in point of care settings to accelerate malaria diagnosis replicating human thought processes.

Moreover, we demonstrated the ability of neural networks in generating synthetic images which are hard to be distinguished from real ones by experienced microscopists. This paves the way to the use of generated images to normalize and translate data from different centers or stainings, offering novel valuable solutions against malaria.

## Author’s contributions

AC designed the study and supervised the project and finalized the manuscript. RTCR conducted all experiments, reported analysis and wrote the intial version of the paper. EOG helped during the analysis and writing of the paper.

## Acknowledgements

We would like to thank Ekene Kwabena Nwaefuna of Ghana Atomic Energy Commission for participaitng to the validation test between real and synthetic images.

